# Atlas of tissue-specific and tissue-preferential gene expression in ecologically and economically significant conifer *Pinus sylvestris*

**DOI:** 10.1101/2020.10.22.350058

**Authors:** Sandra Cervantes, Jaana Vuosku, Dorota Paczesniak, Tanja Pyhäjärvi

## Abstract

Despite their ecological and economical importance, conifers genomic resources are limited, mainly due to the large size and complexity of their genomes. Additionally, the available genomic resources lack complete structural and functional annotation. Transcriptomic resources have been commonly used to compensate for these deficiencies, though for most conifer species they are limited to a small number of tissues, or capture only a fraction of the genes present in the genome.

Here we provide an atlas of gene expression patterns for conifer *Pinus sylvestris* across five tissues: embryo, megagametophyte, needle, phloem, and vegetative bud. We used a wide range of tissues and focused our analyses on the expression profiles of genes at tissue level. We provide comprehensive information of the per-tissue normalized expression level, indication of tissue preferential upregulation and tissue-specificity of expression. We identified a total of 48,001 tissue preferentially upregulated and tissue specifically expressed genes, of which 28% have annotation in the Swiss-Prot database. Even though most of the putative genes identified do not have functional information in current biological databases, the tissue-specific patterns discovered provide valuable information about their potential functions for further studies, as for example in the areas of plant physiology, population genetics, and genomics in general. As we provide information on tissue specificity at both diploid and haploid life stages, our data will also contribute to the understanding of evolutionary rates of different tissue types and ploidy levels.

## Introduction

Conifers, a clade within the gymnosperms, represent a group of plants with significant economic and ecological relevance (Farjon 2008). Several coniferous trees are among the most important sources of wood and timber, as for example *Pinus* and *Picea* (San-Miguel-Ayanz et al. 2016; 2020). Conifers dominate boreal forests worldwide and can form large forested areas hosting a variety of ecosystems. Furthermore, conifer forests are one of the major ecosystem services providers and they are crucial for carbon sequestration (Bonan et al. 1995; DeAngelis 2008; San- Miguel-Ayanz et al. 2016; Boonstra et al. 2016). Despite their importance, genomic resources for conifers, and gymnosperms in general, lag behind in availability compared to angiosperms. Although several contributions have been made recently to fill this gap (Nystedt et al. 2013; Birol et al. 2013; Zimin et al. 2014; Stevens et al. 2016; Mosca et al. 2019), conifer genome annotation remains a challenge, with both structural and functional annotations being far from perfect (Wegrzyn et al. 2014; Cañas et al. 2019). Conifer genomics resources are limited due to the large size of their genomes, ranging from 8 to 70 Gbp (Zonneveld 2012) and to the large number of repetitive elements (approximately 80%) within them (Nystedt et al. 2013; Neale et al. 2014; De La Torre et al. 2020). Proper and complete annotation of the conifer genomes has also been complicated by the presence of long introns (Nystedt et al. 2013; Wegrzyn et al. 2014), which prevents the routine use of common annotation software. Moreover, analyses of ortholog genes across different species indicate that there are several gene groups which are unique to conifers or conifer species specific, with no well-defined homologs in any of the angiosperm plant models (Nystedt et al. 2013; Wegrzyn et al. 2014; Neale et al. 2014; Baker et al. 2018).

Transcriptomic resources have been particularly important for research in conifers and other non-model species, as a strategy to compensate for the challenges associated with efficient genome assembly and annotation (Cañas et al. 2019; Wegrzyn et al. 2020). As the biological functions can not be directly inferred from nucleotide sequences, reference transcriptomes and gene expression studies are useful in the identification and annotation of genes (Raherison et al. 2012; Wegrzyn et al. 2014; Merino et al. 2016; Little et al. 2016; Canas et al. 2017). Transcriptome information can also be used in conifers that lack reference genomes, as this information can be used in the design of reduced genome representation targets (Rellstab et al. 2019; Tyrmi et al. 2020). In addition to this, RNA-seq analyses allow the identification of expression patterns and expression levels, which are essential components of evolutionary genomics studies. For example, selective constraints in genes can be inferred from their expression patterns, as both breadth and expression level are known determinants of evolutionary rates (Wright et al. 2004; Slotte et al. 2011). Selective constraints are also expected to differ between haploid and diploid tissues which differ in the relative rate of expression, as tissue specificity and ploidy has potentially drastic effects on the dynamics of e.g. purifying selection (Otto et al. 2015).

Here we give a first glimpse of the expression patterns of tissue preferentially upregulated (PUR) and tissue specifically expressed genes across five tissues (embryo, megagametophyte, needle, phloem, and vegetative bud) of *Pinus sylvestris. P. sylvestris* is a widely distributed conifer of large economic and ecological importance in Northern Eurasia (Pyhäjärvi et al. 2020). *P. sylvestris* is one of the main sources of timber and raw material for the pulp and paper industry in Europe and is a dominant species in boreal forests, with an estimated coverage area of 145 millions hectares (Pyhäjärvi et al. 2020). *P. sylvestris* is also a suitable model to answer evolutionary and genetic questions, especially regarding gymnosperm reproductive biology, its evolution and genetic consequences. For example, in conifers the maternal nuclear haplotype of an embryo is identical to the megagametophyte’s nuclear haplotype (Williams 2008), which makes it possible to separate expression of paternal and maternal haplotypes and alleles in the embryo (Verta et al. 2016).

Despite its importance and potential, *P. sylvestris* still lacks a reference genome, and currently there are limited genomic resources for this species (see however (Wachowiak et al. 2015; Merino et al. 2016; Li et al. 2017; Höllbacher et al. 2017; Ojeda et al. 2019; Perry et al. 2020)). To date, the few transcriptomic studies of *P. sylvestris* have been based on a small number of tissue types such as needles or seed tissues (Wachowiak et al. 2015; Merino et al. 2016). Identification of tissue preferentially upregulated and tissue specific genes is relevant because 1) understanding the different patterns of expression across different kinds of tissues can aid to elucidate the organization of transcriptomes (Raherison et al. 2012). 2) Knowing the different profiles of expression across tissues can set the ground for evolutionary analysis, as it is known from studies in mammals and angiosperms that the evolution of gene expression differs across tissues or organs (Brawand et al. 2011; Yang and Wang 2013). Ultimately this knowledge will help to gain a deeper understanding of the determinants and main factors that affect the rate of adaptive evolution and the dynamics at the genome level.

In this study we 1) provide a comparative transcriptomic resource for *P. sylvestris* describing the expression level in five different tissues, 2) identify genes that are tissue preferentially upregulated and tissue specifically expressed in each of the five tissues, 3) provide quantitative measures of tissue-specific expression for each gene per tissue combination, and 4) conduct gene ontology enrichment analysis for each tissue type. Our results are important for future studies in comparative conifer genomics, plant physiology, population genetic analyses, evolutionary genetic studies, further gene expression analyses, and aid in the annotation of present and forthcoming conifer genome sequences.

## Materials and Methods

### Plant material

During the growing season of 2016 (May 26th-27th), we sampled needles, phloem, vegetative buds (called tissues for brevity, but see results and discussion section), and seeds from six non-related adult *Pinus sylvestris* trees growing at the Punkaharju Intensively Studied Site (ISS) (http://www.evoltree.eu/index.php/intensive-study-sites) in Southern Finland, resulting in total of 30 samples (Table S1). The site is a natural forest stand dedicated to forestry research conducted by Natural Forest Research Institute Finland (LUKE). Samples were collected in collaboration with LUKE [33] that has an agreement on the forest research use with the owner Metsähallitus (The Finnish Forest Administration). *P. sylvestris* in Finland is not endangered and Finland does not regulate its genetic resources under Nagoya Protocol so CITES was not applied and prior informed consent was not needed. The samples were collected and identified by Tanja Pyhäjärvi. There is no voucher available for the specimen. The same plant material and sequenced libraries used in this work, have been used previously to assemble multiple reference transcriptomes of *P. sylvestris (Ojeda et al. 2019)* and a more detailed description of the plant material and RNA extraction procedure is described by Ojeda et al. (2019). Needle, phloem, and bud samples were stored immediately in RNAlater (Thermo Fisher Scientific) or frozen *in situ* and transported to the storage on dry ice. Samples were stored at −80°C (samples in dry ice) or −20°C (samples in RNAlater) until RNA extraction. We obtained megagametophyte and embryo tissues by dissecting mature seeds collected from the same mother trees from which the vegetative tissues were obtained. Seeds were stored in the dark at 4°C until germination was induced by exposure to moisture and continuous light for 48 h.

### RNA isolation, library preparation and sequencing

We extracted total RNA from needle, bud, and phloem using the Spectrum Plant Total RNA Kit (Protocol B, Sigma). Total RNA extraction was followed by mRNA capture with the NEBNext® Poly(A) mRNA Magnetic Isolation Module (New England Biolabs Inc.). For embryo and megagametophyte, mRNA was directly extracted from the whole tissues with Dynabeads mRNA Direct Micro Kit (Thermo Fisher Scientific) according to manufacturer’s protocol, except for using 200 μl of lysis buffer. RNA concentration was quantified with Qubit RNA HS Assay kit (Thermo Fisher Scientific). We prepared a total of 30 libraries using the NEBNext Ultra Directional RNA Library Prep Kit for Illumina (New England Biolabs Inc.). We selected an insert size of 300 bp by using a fragmentation time of 5-12 min, followed by size selection with 40-45 μl *I* 20 μl AMPure XP beads (Agencourt). Libraries were single indexed with NEBNext Multiplex Oligos Set 1 for Illumina, and finally enriched with 12-15 PCR cycles. We quantified the libraries and visually checked the fragment size distribution before sequencing. We used paired-end (2 x 150 bp) and sequenced five pools of 6 to 12 libraries on an Illumina NextSeq 500 at the Biocenter Oulu Sequencing Centre (Oulu, Finland).

### Transcript quantification

We used trimmed reads (BioProject PRJNA531617) as input for transcript quantification, using the Trinity_guided_ [8] as a reference transcriptome Data S1). We followed the Trinity Post- Transcriptome Assembly Downstream Analyses pipeline (Trinity v. 2.6.6) (trinityrnaseq; Haas et al. 2013) to generate quantification files at isoform level, and raw counts and normalized count matrices at putative gene level (hereafter referred as gene level matrices). We first obtained transcript abundance independently for each of the six individuals in each one of the five tissues. This was done by pseudo-aligning the RNA-seq reads to the transcriptome reference with Salmon 0.9.1 (Patro et al. 2017) using the --SS_lib_type (strand specific) and --trinity_mode options. The -- trinity_mode option allowed the estimation of counts from isoforms to generate counts at a putative gene level during the count matrix generation step. Before any further analysis, we checked for the presence of possible contaminants by searching contigs that had hits to the keywords ‘alveolata’, ‘metazoa’, ‘fungi’, ‘bacteria’, and ‘archaea’. We search for exact matches to these keywords from the results of a translated blast (BLASTX) of the transcriptome annotation file (Ojeda et al. 2019; Ojeda 2020). We then combined our list of putative contaminants with the contaminants and organelles contigs lists reported in Ojeda et al. (2019), and excluded them from the isoform quantification files and the gene_trans_map. Contaminants were removed after the pseudo-aligning stage to avoid the false mapping of contaminant reads to non-contaminant contigs in the reference transcriptome.

### Abundance matrices construction

We built three count matrices with the Trinity pipeline (abundance_estimates_to_matrix.pl) at the gene level based on the cleaned independent transcript quantification. First, we generated a gene level raw counts matrix (Table S2), which was then used to construct a transcript per million length normalized gene count matrix (TPM escalated matrix) (Table S3). The TPM escalated matrix accounts for differences in isoform lengths that otherwise could inflate FDR due to differential transcript usage (Soneson et al. 2015). Finally, the TPM escalated matrix was used to construct a gene counts matrix normalized using the Trimmed Mean of M values (TMM) method (Table S4), which accounts for differences in the distribution of transcript expression that could lead to an increase in false positive rates, and decrease the power to detect truly differentially expressed genes (Robinson and Oshlack 2010). Before doing the differential expression analyses and the estimation of tissue specificity, we evaluated the quality of our samples by doing a principal component analysis (PCA) and a Pearson correlation matrix using the gene raw count matrix, according to the Trinity QC samples and biological replicates pipeline (trinityrnaseq). The intention of these analyses was to look for the presence of batch effects or sample outliers, and to verify that biological replicates clustered within each tissue type and not among sampled individuals.

### Differential expression analysis and identification of tissue preferentially upregulated genes

Differentially expressed genes (DEG) and PUR genes were identified using the Trinity Differential Expression and Sample-Specific Expression pipelines (trinityrnaseq; Bryant et al. 2017). Briefly, we first identified DEG using the gene raw counts matrix with edgeR 3.28.0 (Robinson et al. 2010; McCarthy et al. 2012). The differential expression analysis was based on pairwise comparisons of each of the 5 tissues (10 pairs), using the six samples per tissue as biological replicates.

For each pair of DEG identified we obtained their associated false discovery rate (FDR), and then we used this information combined with the normalized counts of the TMM matrix to identify the PUR genes in each of the five tissues. We obtained a normalized mean value of expression for each tissue by averaging and log 2 transforming the counts for each gene across the six replicates for each tissue on the TMM gene matrix. Each DEG with a maximum FDR of 0.05 for differential expression and with positive logFC of the log2 transformed gene counts in the TMM matrix was classified as PUR. A summary of pairwise expression differences between tissues based on the logFC of the log2 transformed gene counts in the TMM matrix is provided in Data S2.

### Tissue-specific expression

As an alternative approach to quantitatively assess the tissue-specific expression of the genes we calculated the τ index based on the TMM gene counts matrix. The τ index ranges between 0 for widely expressed genes, and 1 for exclusively tissue-specific genes (Yanai et al. 2005). As the τ index considers tissue specificity independently of the level of expression, we set as “not expressed” genes with expression values < 1 from our TMM matrix in order to exclude genes with low support for true expression and low signal to noise ratio. To do this, we first log2 transformed the matrix in order to normalize the distribution of the expression values. We set all negative values in the matrix to zero, as this represented values < 1 before log2 transformation. We excluded contigs that had no expression values or that had expression in just one out of the 30 samples.

Then, the τ index was computed separately for each gene across all tissues and replicates according to the following equation (Yanai et al. 2005; Kryuchkova-Mostacci and Robinson-Rechavi 2017):

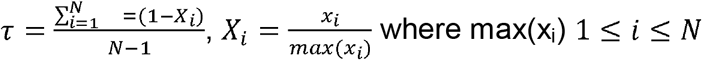

Where N represents the number of tissues, x_i_ is the mean expression in tissue *i* and X_i_ is the expression level in tissue *i* normalized by the maximum mean expression among all tissues (severinEvo).

### Singular enrichment analysis

To further characterize the gene expression in the five tissues, we identified the biological pathways for both tissue-specific and tissue preferentially upregulated gene sets with independent singular enrichment analysis (SEA) (Huang et al. 2009; Du et al. 2010). First, we retrieved the UniProt IDs corresponding to our putative genes from the blastx field from our reference annotation file (Ojeda et al. 2019). Then we uploaded the list of UniProt IDs to the uniprot retrieve/ID mapping tool (https://www.uniprot.org/uploadlists/) and restricted the result to GO terms only. We repeated this procedure with the genes used as a background list for the SEA: all the contigs in the gene raw counts matrix for the PUR genes (Data S3), and all the contigs in the filtered TMM matrix in the case of the tissue-specific genes (Data S4).

Of the 715,398 putative genes in the raw counts matrix used for the differential expression analysis, 17,227 have a unique UniProt ID and represent 108,947 GO terms. The background list for the tissue-specific genes data set consisted of 177,075 contigs of which 14,079 have a unique annotation and represent 90,198 GO terms. For both data sets only uniquely annotated genes and their corresponding GO terms (Data S5-S14) were used for running the singular enrichment analyses to avoid inflating the number of GO terms falsely, and creating a bias in the analysis.

We used the GO terms along the UniProt IDs as input for the SEA using the agriGO (http://systemsbiology.cau.edu.cn/agriGOv2/index.php) platform (Du et al. 2010; Tian et al. 2017). We used the custom background list option, applied a hypergeometric test as statistical test method with a minimum of 5 mapping entries per term, and Hochberg FDR as multi-test adjustment method with a significance level of 0.05.

## Results and discussion

### Transcript quantification and abundance matrices construction

We mapped a total of 707,063,773 trimmed and adapter removed reads from five different tissues (embryo, megagametophyte, needle, phloem, and vegetative bud) and six biological replicates (six different genotypes) per tissue type to *P. sylvestris* TRINITYguided transcriptome (Ojeda et al. 2019). On average 23,568,792 reads originated from each tissue, ranging from 29,591,629 reads for needle to 20,469,80 reads for phloem. On average 76% of the reads per replicate were successfully mapped to the reference (Table S5). After mapping 1,307,500 contigs had aligned reads at the isoform level. Of those, 120,040 contigs were removed from the downstream analyses as they were identified as contaminants (Data S15). The final set consisted of 1,187,460 contigs at isoform level and were used to construct raw counts and normalized matrices at gene level for downstream analyses (see Materials and Methods section). The total number of putative genes with expression signal in the gene level matrices was 715,398, much higher than the number of annotated genes in any conifer (Nystedt et al. 2013; Neale et al. 2014; Gonzalez-Ibeas et al. 2016).

This magnitude, albeit probably an overestimate, is typical to transcriptome studies (Little et al. 2016). This is likely a result of single genes being present in multiple fragments, isoforms split into multiple genes, and different alleles originating from heterozygous material identified as separate genes during assembly and classification as genes by Trinity (Ojeda et al. 2019). However, part of the genes originate from gene families and since clustering similar genes is possible in downstream analysis, we chose to err on the side of potentially over splitting the genes rather than imperfectly clustering similar transcripts as a single gene, as over clustering will inherently lead to loss of information. We believe that providing expression data with minimum clustering will be most versatile for later use of the transcriptome and expression data in genome annotations and other studies.

### Quality assessment of biological replicates

As we used different genotypes as biological replicates, we first verified that the replicates clustered by tissue type and not by genotype, and checked for the presence of potential outliers in the dataset. We used the raw counts matrix data (Table S2), a principal component analysis (PCA) and a Pearson correlation to verify this. The PCA separated the tissue samples into five distinct clusters without any overlap, indicating that among-tissue variation is the main factor of among-sample variation (Figure 1). Hence, our approach captures the differentiating gene expression profiles of the five tissues. In the PCA, the seed-derived megagametophyte and embryo samples clustered closest to each other, suggesting similarity in their gene expression profiles. Also phloem and bud samples clustered close to each other, whereas needle samples showed the most unique gene expression profile. In the hierarchical clustering analysis, based on the correlations of gene expression profiles, the differences among tissues are relatively shallow. But, similarly to the PCA, all replicates are clustered according to their tissue type and not according to their genotypes, corroborating the PCA results (Figure S1).

**Figure 1.**
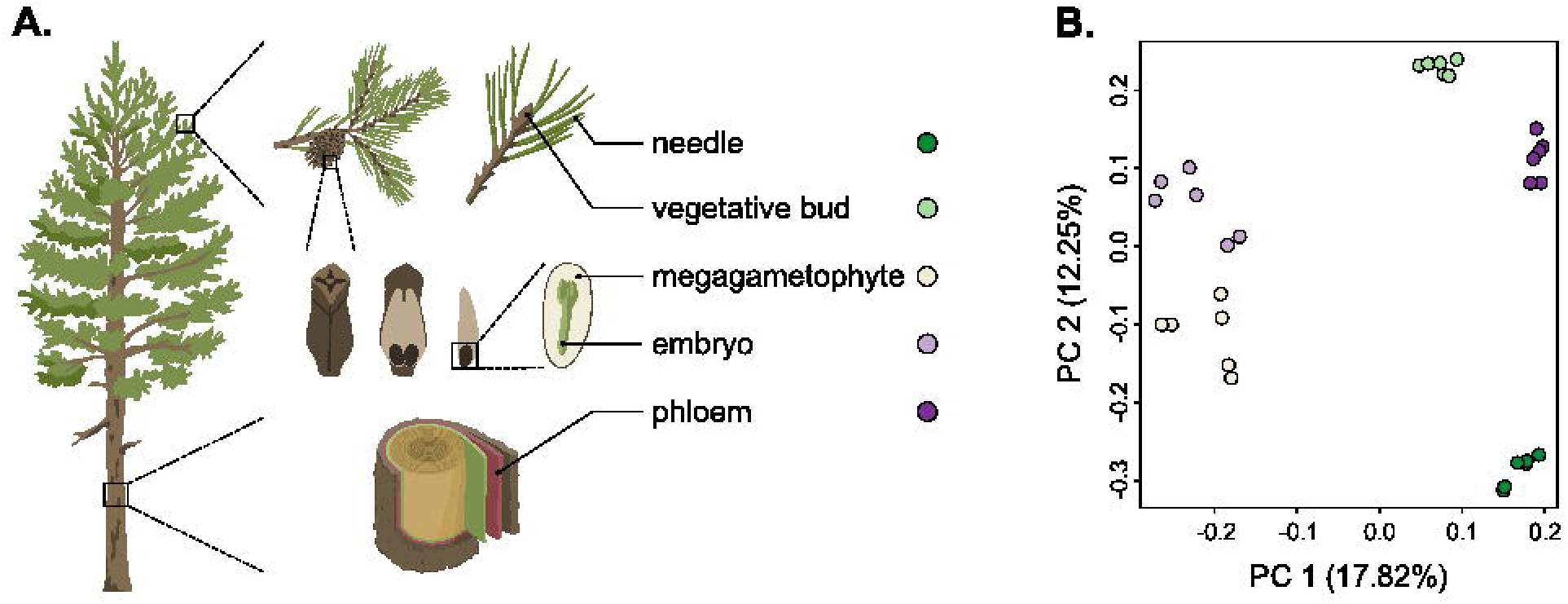
**A)** Schematic representation of the five tissues used in the transcriptome profiling of *Pinus sylvestris:* needle, vegetative bud, megagametophyte, embryo and phloem. **B)** Scatterplot of the first two axes of the principal component analysis (PCA). Tissue types are denoted by colors

### Tissue preferentially upregulated and tissue-specific gene expression

We defined a gene as tissue PUR when there was a significant log fold change in the expression value compared to the other tissues. To identify tissue PUR we first did a differential expression (DE) analysis. For this we included all the genes in the raw count matrix (Table S2). We decided not to apply any minimum number of counts per gene as a filtering threshold to run the analysis, as we later applied a 5% false discovery rate (FDR) threshold for the identification of PUR genes. Out of the 715,398 genes initially included in the DE analysis, 198,413 genes had a maximum 5% FDR for differential expression and were further included in the analysis to identify PUR genes. We identified a total of 48,001 genes with tissue preferential expression, and out of the five tissues needle has the highest number of PUR genes (Table 1)

**Table 1.**
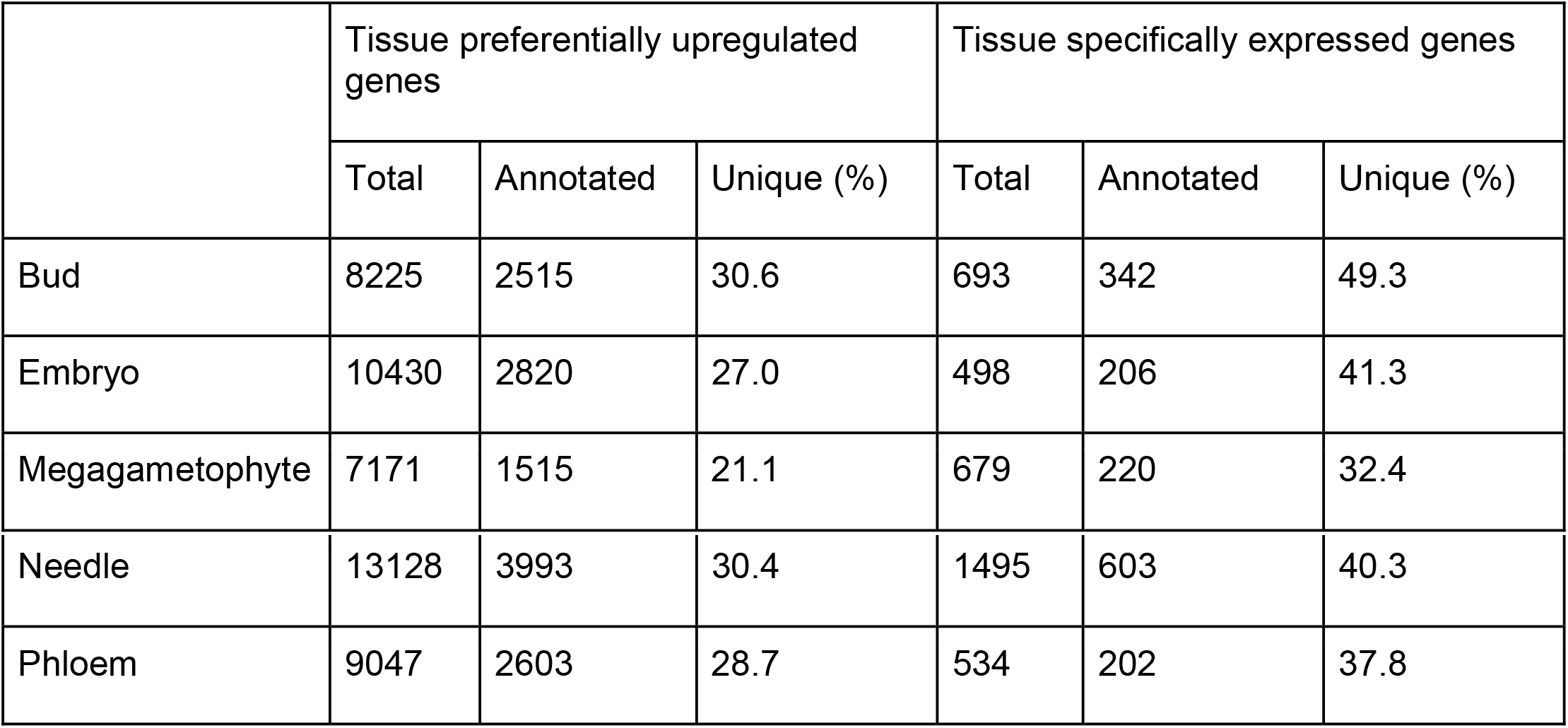
Number of genes identified as tissue preferentially upregulated and tissue-specific in five *P. sylvestris* tissues. The percentage of unique UniProtKB identifiers is also shown.

|  | Tissue preferentially upregulated genes |  |  | Tissue specifically expressed genes |  |  |
| --- | --- | --- | --- | --- | --- | --- |
|  | Total | Annotated | Unique (%) | Total | Annotated | Unique (%) |
| Bud | 8225 | 2515 | 30.6 | 693 | 342 | 49.3 |
| Embryo | 10430 | 2820 | 27.0 | 498 | 206 | 41.3 |
| Megagametophyte | 7171 | 1515 | 21.1 | 679 | 220 | 32.4 |
| Needle | 13128 | 3993 | 30.4 | 1495 | 603 | 40.3 |
| Phloem | 9047 | 2603 | 28.7 | 534 | 202 | 37.8 |

Quantification of tissue specificity allows a powerful statistical analysis of correlation between tissue-specific expression and e.g. evolutionary rate or other dependent or explanatory variables and factors. We identified the tissue specifically expressed genes by calculating the τ score per gene. The score ranges from zero to one, with a zero given to genes expressed in all tissues and one given to completely tissue specific genes. For this analysis we retained a set of 177,075 genes (Table S6) after applying the filtering criteria described in Methods. We considered a gene as tissue specifically expressed only if its ⊺=1. We identified a total of 3,899 genes with a tissue-specific pattern of expression, similarly the PUR analysis results, needle has the highest number of tissuespecific genes (Table 1). To obtain the annotation of the genes identified as tissue PUR and tissue specific, we retrieved the corresponding UniProtKB identifiers (Ojeda 2020) from the Trinotate for the 715,398 putative genes in the TMM count matrix, out of which 97,435 (14 %) had a Swiss-Prot (Bairoch and Apweiler 2000) protein match based on BLASTX (Ojeda et al. 2019). Most of the Swiss-Prot annotations (67%) originated from *Arabidopsis thaliana* (65,214 genes). Other common annotation sources were *Nicotiana tabacum* (9,794; 10%) and *Oryza sativa* (8,946; 9%). Only 1663 genes (1.7%) had an annotation to other *Pinus* species, of which 177 (10.6%) were hits to *P. sylvestris,* and 608 (36.5%) genes had Swiss-Prot annotation to *Picea.* Note that Swiss-Prot is a manually curated database that does not currently have a comprehensive set of annotated gymnosperm proteins and therefore the best matches are often obtained from the model plants such as *A. thaliana.* A proportion of our putative genes share the same gene identifier (annotation) (Table 1). This probably reflects the incomplete collapse of different isoforms in the assembled transcriptome used as reference, or the presence of gene families (Wegrzyn et al. 2014). Also, a high number of the genes identified as PUR or tissue specific lack annotation altogether, which is not surprising as genes with higher tissue-specific expression have less conserved sequences and are less likely to find orthologs among other species (Lemos et al. 2005; Raherison et al. 2012). A summary of the 715,398 genes indicating their normalized expression level (TMM), τ score, tissue specificity status, PUR status, and annotation can be found in the Supplementary information (Table S7).

Cursory inspection of annotations of highly expressed tissue PUR and tissue-specific genes are congruent with some of the already known functions of the tissues. These results confirm that our analyses capture biologically meaningful characteristics of the tissues. For example in megagametophytes, enzymes related to seed storage lipid mobilization and germination were upregulated and specifically expressed. Similarly, in needles, several chlorophyll a-b binding proteins are upregulated. In embryo, multiple ribosomal proteins and other proteins indicating active protein synthesis were upregulated. In vegetative buds, expression of genes involved in defense against insect attack, like (-)-alpha-pinene synthase and dirigent (Ralph et al. 2006) that take part in oleoresin synthesis, were highly expressed and specific to this tissue. In phloem, the two genes annotated as metallothionein-like protein EMB30, an aquaporin and a thioredoxin-like protein were highly expressed, similarly to *Quercus suber* phellem (cork) where metallothionein reacts to oxidative stress (Mir et al. 2004) or in *Pinus taeda* xylem where the same proteins were among the most highly expressed genes (Lorenz and Dean 2002).

Among the five tissues analysed, the needle had the highest number of genes with tissue-specific expression and embryo the lowest (Table 1). Except for two genes, one in megagametophyte and one in needle, all the genes with tissue-specific expression were also among the PUR genes. However, as tissue specificity does not require a high expression level, genes with τ score equal to one are not necessarily the most upregulated genes in their respective tissues. Comparison of our findings to other studies is not straightforward as there are very few transcriptomic studies in *P. sylvestris.* But in comparison to a previous study (Merino et al. 2016), where they focus on the comparison between megagametophyte and embryo tissues at different developmental stages, we identified less megagametophyte and embryo specifically expressed genes. One of the reasons for this difference could be that the identification of unique genes in the previous study [20] was based only on the comparison between embryo and megagametophyte tissues. As the identification of tissue specific genes is contingent to the number of tissues used for the analysis, it is expected that the higher the number of tissues used in the comparison, the lower the number of tissue specific genes that will be identified. In contrast, we found a higher number of tissue specific genes in embryo, bud, and needle compared to a previous study in conifers (Raherison et al. 2012), where several tissue types were used. One notable difference between this (Raherison et al. 2012) and ours was the higher number of tissue-specific genes for megagametophyte found in *P. glauca.* This analysis (Raherison et al. 2012) found the highest number of unique genes in the megagametophyte in comparison to other tissues analyzed. The low number of megagametophyte specific genes identified in our study could be due to the use of mature embryos as starting material. Previous research suggests that the number of unique transcripts in the megagametophyte varies during the developmental stages of embryogenesis (Merino et al. 2016).

One caveat of our analyses is that, unlike other studies, we did not use microdissection in order to obtain the tissue samples (Canas et al. 2017). Hence, some of the “tissues” are a mix of tissue types. Needles, for example, include several tissues (phloem among them) (Pongrac et al. 2019), and mature embryos contain the shoot and root meristems as well as cotyledons (Singh 1978). In contrast, the mature megagametophyte is a quite uniform storage tissue consisting of cells packed with starch protein and lipids (Simola 1974; Vuosku et al. 2015). Another limitation of the dataset is that it represents only one point in time and space, although gene expression is a dynamic process and quantitative and qualitative variations exist over spatial and temporal scales. Instead of sampling across several developmental stages or across a spatial gradient our dataset represents a wider set of tissues, which increases the power to identify tissue PUR and tissue specifically expressed genes. The added value of the dataset lies in the unexpected functions and connections discovered among biological pathways and genes with previously unidentified signals of tissue-specificity or up-regulation.

### Functional characterization of tissue preferentially upregulated and tissue-specific genes

GO enrichment analysis allows the identification of gene functions enriched with certain functional roles. The number of enriched functions was of the same magnitude across tissue types, ranging from 253 to 452 for PUR genes and from 58 to 169 for tissue-specific genes (Table S8-S17). The total number of GO terms and the number of significant enriched terms per tissue are shown in Table 2, a summary of the most highly expressed genes per tissue, and the most significantly enriched biological processes is shown in Figure 2. Most of the genes (86%) with expression signals in our study lacked annotation from the Trinotate pipeline. Thus, they did not contribute to functional analysis or GO enrichment results.

**Table 2.** Total number and number of significant GO terms and percentage of enriched terms in *P. sylvestris* tissues.

|  | Tissue preferentially upregulated genes |  |  | Tissue-specific genes |  |  |
| --- | --- | --- | --- | --- | --- | --- |
|  | Total | Significant | Percentage (%) | Total | Significant | Percentage (%) |
| Bud | 15681 | 452 | 2.9 | 2019 | 137 | 6.7 |
| Embryo | 17461 | 253 | 1.4 | 1178 | 75 | 6.4 |
| Megagametophyte | 9690 | 306 | 3.1 | 1363 | 111 | 8.1 |
| Needle | 25295 | 401 | 1.6 | 3818 | 169 | 4.3 |
| Phloem | 16371 | 422 | 2.6 | 1249 | 58 | 4.6 |

The complete lists of gene identifiers and their corresponding GO terms per tissue and per each set of genes Data S5-S14), along with tables with the results of the SEA showing each GO terms, its p-value, and FDR (Table S8-S17) are provided in supplementary information.

**Figure 2.**
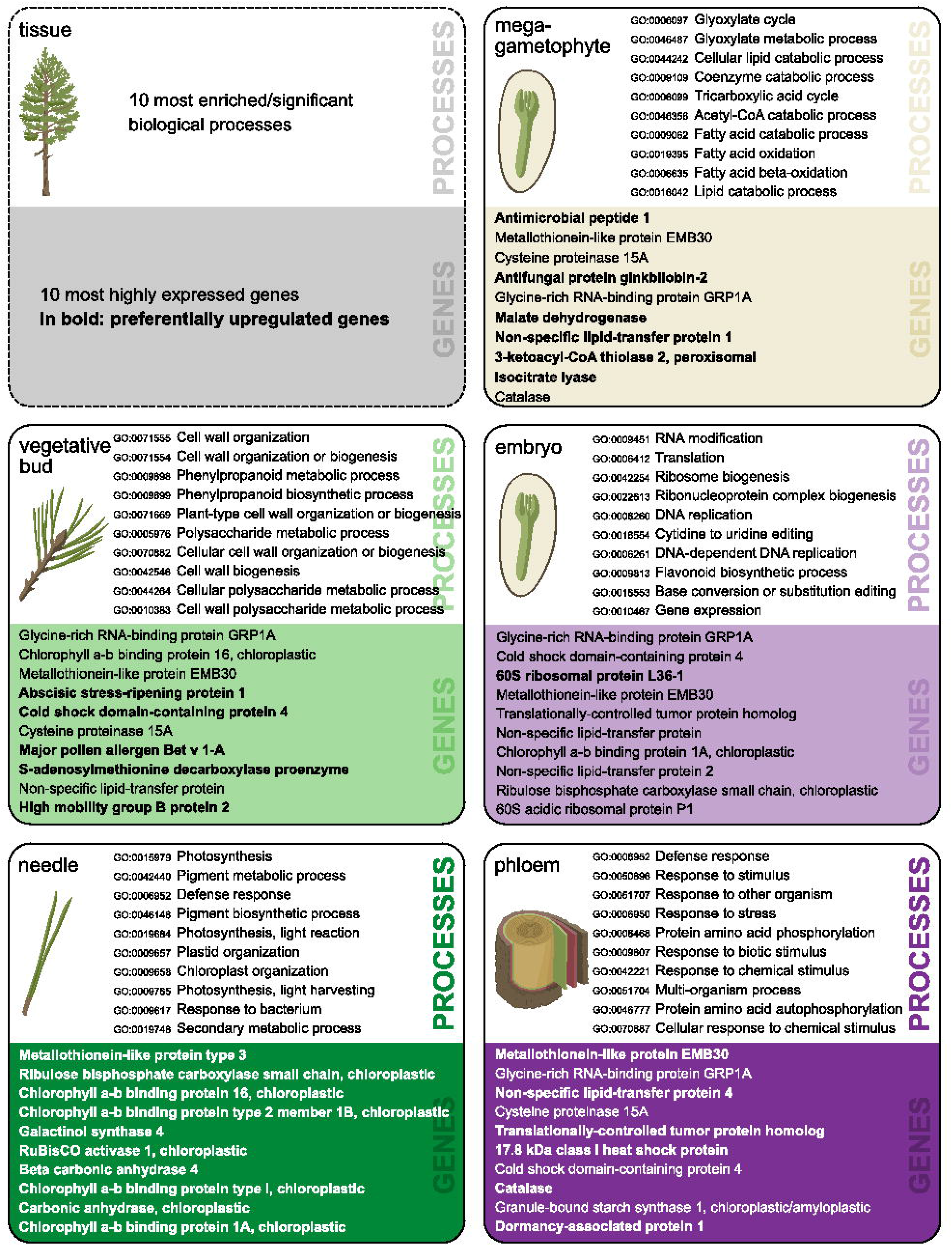
Ten most significantly enriched biological processes (with corresponding GO-term IDs) and ten most highly expressed annotated genes in each of the five tissues. Genes preferentially upregulated (PUR) in a given tissue are in bold

In needles the significant GO terms reflected the exposure of trees to various stresses and interactions with other organisms, whereas in embryos, buds and the phloem significant GO terms were mainly connected to different development-related processes. In needles the enriched biological process GO terms among tissue-specific genes were immune response (GO:0006955) as well as response to stress (GO:0006950) and other organisms (GO:0051707) such as oomycetes (GO:0002229), bacteria (GO:0042742) and fungi (GO:0009817). Moreover, terpene synthase activity (GO:0010333), which may play a key role in the defense against herbivores (Achotegui- Castells et al. 2013), was an enriched molecular function among tissue-specific genes in needles, but also in embryos and vegetative buds. For example, reactive oxygen species (ROS) related biological processes (GO:0006800, GO:0042743, GO:0034614) and molecular functions (GO:0004601, GO:0004364) were enriched among the tissue-specific genes in embryos consistent with an active ROS protection in developing tissues. In the phloem, a special differentiation process, syncytium formation (GO:0006949), indicating the interconnection of phloem sieve elements to generate a transport route (Geldner 2014) was an enriched biological process among the tissue specific genes.

### Megagametophyte-specific genes have crucial functions in seed germination and energy conversion

Gymnosperms are characterized by the haploid female gametophyte tissue, the megagametophyte, which surrounds the embryo in developing and mature seeds. The megagametophyte can be considered a functional homolog of the endosperm in angiosperms due to its role as a nourishing tissue (King and Gifford 1997; Costa et al. 2004). However, the megagametophyte develops from a haploid megaspore before the fertilization (Singh 1978) and is therefore entirely maternally inherited unlike the diploid or triploid endosperms of biparental origin (Williams and Friedman 2002; Baroux et al. 2002). To give an example of the potential uses of the dataset, we provide a more detailed description of the megagametophyte expression profile, but leave the in-depth analysis of the other tissues for later investigations.

Among highly expressed and up-regulated genes in the megagametophyte were malate synthase (EC 2.3.3.9) and isocitrate lyase (EC 4.1.3.1) that are essential in glyoxylate cycle converting lipids into carbohydrates in seeds (Ching 1970), as well as other glyoxysomal proteins like Acetyl-CoA acyltransferase (EC 2.3.1.16), ABC transporter and peroxisomal fatty acid beta-oxidation multifunctional protein AIM1 (Graham 2008). Seed storage related genes such as 2S seed storage-like protein, 11S globulin seed storage protein 2 and 13S globulin basic chain and some isocitrate lyase copies were completely megagametophyte-specific (⊺=1). Antimicrobial and antifungal protein coding genes were the most highly expressed among annotated megagametophyte-upregulated genes.

The enriched GO terms of biological processes and molecular functions in the megagametophyte tissue-specific genes included seed germination and the mobilization of nutrient reserves. Nutrient reservoir activity (GO:0045735) indicated the mobilization of energy sources from the megagametophyte for seed germination and early seedling growth, as well as lipid catabolic processes (e.g. GO: 0016042, GO:0044242). Malate dehydrogenase activity (GO:0016615) and heme binding (GO:0020037), which mostly originated from the cytochrome P450 enzymes containing heme cofactors (Xu et al. 2015), reflected the resume of active metabolism. Also, response to ROS (GO:0034614) and antioxidant activity (GO:0016209) suggested active metabolism and signaling. ROS are natural by-products of metabolism and may be detrimental to seed viability because they can cause oxidative stress. However, in the seed ROS also work as signals which underpin the breaking of dormancy and provide protection against pathogens (Jeevan Kumar et al. 2015). Megagametophyte cells showed responses to hormone stimulus (GO: 0032870) and the function of hormone-mediated signaling pathways (GO:0009755) including abscisic acid (GO:0009738), auxin (GO:0009734) and ethylene (GO:0009873) which also belong to the molecular networks regulating seed dormancy and germination (Seo et al. 2009; Guangwu and Xuwen 2014; Miransari and Smith 2014; Shu et al. 2016). Cellulose biosynthetic process (GO:0030244) and primary cell wall biogenesis (GO:0009833) suggest that cell walls in the megagametophyte may participate in water retention and give mechanical support to the germinating embryo (Otegui). Similarly to previous findings in *P. sylvestris* (Merino et al. 2016) megagametophytes, we found enrichment for processes involved in the response to chemical and endogenous stimuli (GO:0042221, GO:0071495). Merino et al. (2016) (Merino et al. 2016) suggested that the megagametophyte could also be involved in the regulation of the embryo development through the induction of signaling pathways triggered by sensing environmental signals in a similar way the angiosperms’ endosperm does (Yan et al. 2014). Altogether, our findings show that the megagametophyte is not just a reserve nutrition for the germinating embryo, but a metabolically active tissue contributing in multiple ways to seed germination and, thus, underline the importance of the haploid stage in *P. sylvestris* life cycle.

Several enzymes widely used in allozyme-based population genetic studies ((Szmidt and Muona 1989) and references therein) such as aconitate hydratase (EC 4.2.1.3), malate dehydrogenase (EC 1.1.1.37) and aspartate aminotransferase (EC 2.6.1.1) were megagametophyte-specific and among the top 50 expressed genes in the tissue. As they may be more prone to natural selection against recessive deleterious variants when expressed at the haploid stage, early population genetic analyses may have bias in e.g. estimates of the overall genetic diversity based on these loci.

### Conclusions

We provide a widely and interdisciplinary applicable genome-wide atlas of tissue-level transcription patterns based on RNA-seq for economically and ecologically significant coniferous tree *P. sylvestris.* Quantitative data and analysis of expression level, as well as breadth and tissue specificity are provided for 715,398 different putative genes. The mapping and bioinformatic analyses of gene expression are based on the most complete and high-quality reference transcriptome of *P. sylvestris* available to date (Ojeda et al. 2019). Previous transcriptome studies of *P. sylvestris* have concentrated on a narrow set of tissues in each study such as wood (Paasela et al. 2017), embryo (Merino et al. 2016), and needles (Wachowiak et al. 2015; Duarte et al. 2019) or focused on a limited set of genes (Guseva et al. 2020). The present study allows comparison across a wide set of genes expressed in the above-ground parts of adult *P. sylvestris* trees growing in a natural forest.

In addition to genome sequence annotations, we foresee multiple potential uses for the dataset. Level and breadth of gene expression are known to be linked to the evolutionary rate and level of conservation (ref). By combining our data with similar data in other conifers or angiosperms it is possible to study the evolutionary conservation of expression patterns, or the differences in evolutionary rates across tissue-specific expression levels and gain a deeper understanding of the determinants and main factors affecting e.g. rate of adaptive evolution and dynamics at the genome level. The response of trees to a combination of different stresses is unique and cannot be directly extrapolated from studying only single stressors in experimental conditions (Niinemets 2010). The transcriptome resource for adult *P. sylvestris* trees growing under natural conditions, where they are simultaneously exposed to a number of different abiotic and biotic stresses as well as interactions with other organisms, provides a valuable tool also for physiological studies. Finally, un-annotated conifer genes with high expression or tissue specificity can open up whole new research avenues, independent of the previously available knowledge based on angiosperm model plants such as *A. thaliana* and *Populus*.

## Declarations

### Funding

The Academy of Finland grants 287431, 293819, and 319313 to TP. Part of the work was carried out with the support of Biocenter Oulu to SC.

### Competing interest

The authors declare that they have no competing interests.

### Ethics approval

Not applicable

### Consent to participate

Not applicable

### Consent for publication

Not applicable

## Data availability

Clean reads corresponding to each of the five tissues used in the transcript quantification can be found in BioProject PRJNA531617. The Trinotate file used to obtain the gene identifiers for the tissue PUR and tissue-specific genes identified in this work is at https://doi.org/10.6084/m9.figshare.12865997.v1.

Data S1 Trinity_guided_ transcriptome used as reference (contaminants included) is at https://doi.org/10.6084/m9.figshare.12865997.v1

## Supplementary information

**Figure S1.** Heat map showing the correlation between the expression patterns of the five tissues. A=embryo, KS=vegetative bud, M=megagametophyte, N=needle, Ni=phloem.

**Table S1.** Table with information about the geographical location of the trees used in this study.

**Table S2.** Table including the raw counts per genes across the five tissues.

**Table S3.** Table containing the TPM normalized gene counts. A=embryo, KS=vegetative bud, M=megagametophyte, N=needle, Ni=phloem.

**Table S4.** Table containing the TMM normalized unfiltered gene counts. A=embryo, KS=vegetative bud, M=megagametophyte, N=needle, Ni=phloem.

**Table S5.** Table including information about the number of reads mapped to each one of the tissues.

**Table S6.** Table including the TMM normalized and filtered counts used for the identification of genes with tissue-specific expression patterns. A=embryo, KS=vegetative bud, M=megagametophyte, N=needle, Ni=phloem.

**Table S7.** Table including the level of expression, indication of tissue PUR or tissue-specific expression, and annotation for 715,398 genes across the five tissues. Column one indicates gene ID, columns two to six contain the TMM normalized gene counts per tissue, column seven indicates the gene tau score, columns eight to 12 indicate if in which tissue the gene is preferentially upregulated, column 13 indicates the UniProt ID, column 14 indicates the protein name, column 15 indicates the gene name, column 16 indicates the organism from where the annotation was obtained.

**Table S8.** Table containing the results of the SEA for tissue PUR genes expressed in vegetative bud. Column two indicates the ontological process where P=biological processes, F=molecular function, C=cellular component.

**Table S9.** Table containing the results of the SEA for tissue PUR genes expressed in embryo. Column two indicates the ontological process where P=biological processes, F=molecular function, C=cellular component.

**Table S10.** Table containing the results of the SEA for tissue PUR genes expressed in megagametophyte. Column two indicates the ontological process where P=biological processes, F=molecular function, C=cellular component.

**Table S11.** Table containing the results of the SEA for tissue PUR genes expressed in needle. Column two indicates the ontological process where P=biological processes, F=molecular function, C=cellular component.

**Table S12.** Table containing the results of the SEA for tissue PUR genes expressed in phloem. Column two indicates the ontological process where P=biological processes, F=molecular function, C=cellular component.

**Table S13.** Table containing the results of the SEA for genes with tissue-specific expression pattern in vegetative bud. Column two indicates the ontological process where P=biological processes, F=molecular function, C=cellular component.

**Table S14.** Table containing the results of the SEA for genes with tissue-specific expression pattern in embryo. Column two indicates the ontological process where P=biological processes, F=molecular function, C=cellular component.

**Table S15.** Table containing the results of the SEA for genes with tissue-specific expression pattern in megagametophyte. Column two indicates the ontological process where P=biological processes, F=molecular function, C=cellular component.

**Table S16.** Table containing the results of the SEA for genes with tissue-specific expression pattern in needle. Column two indicates the ontological process where P=biological processes, F=molecular function, C=cellular component.

**Table S17.** Table containing the results of the SEA for genes with tissue-specific expression pattern in phloem. Column two indicates the ontological process where P=biological processes, F=molecular function, C=cellular component.

**Data S1.** Trinity_guided_ transcriptome used as reference (contaminants included). At https://doi.org/10.6084/m9.figshare.c.5181788.v1

**Data S2.** Pairwise expression differences between tissues.

**Data S3.** List of genes used as a background list for the SEA of PUR genes.

**Data S4.** List of genes used as a background list for the SEA of tissue-specific genes.

**Data S5.** Genes identified in vegetative bud with PUR expression pattern and their associated GO terms.

**Data S6.** Genes identified in embryo with PUR expression pattern and their associated GO terms.

**Data S7.** Genes identified in megagametophyte with PUR expression pattern and their associated GO terms.

**Data S8.** Genes identified in needle with PUR expression pattern and their associated GO terms.

**Data S9.** Genes identified in phloem with PUR expression pattern and their associated GO terms.

**Data S10.** Genes identified in vegetative bud with tissue-specific expression pattern and their associated GO terms.

**Data S11.** Genes identified in embryo with tissue-specific expression pattern and their associated GO terms.

**Data S12.** Genes identified in megagametophyte with tissue-specific expression pattern and their associated GO terms.

**Data S13.** Genes identified in needle with tissue-specific expression pattern and their associated GO terms.

**Data S14.** Genes identified in phloem with tissue-specific expression pattern and their associated GO terms.

**Data S15.** List of contigs identified as contaminants.

## Authors’ contributions

SC: Main responsibility of the statistical and bioinformatic analyses, main responsibility of writing and editing the manuscript.

JV: Writing parts of the manuscript, editing and commenting the manuscript.

DP: Visualization of the data, editing and commenting the manuscript.

TP: Concept, laboratory work, acquisition of funding, writing and editing the manuscript.

## Acknowledgments

The authors wish to acknowledge CSC - IT Center for Science, Finland, for computational resources.

## Notes

https://doi.org/10.6084/m9.figshare.c.5181788.v1

### Competing Interest Statement

The authors have declared no competing interest.

