## Supplementary figures and images for "Atlas of tissue-specific and tissue-preferential gene expression in ecologically and economically significant conifer *Pinus sylvestris*"

### Fig S1

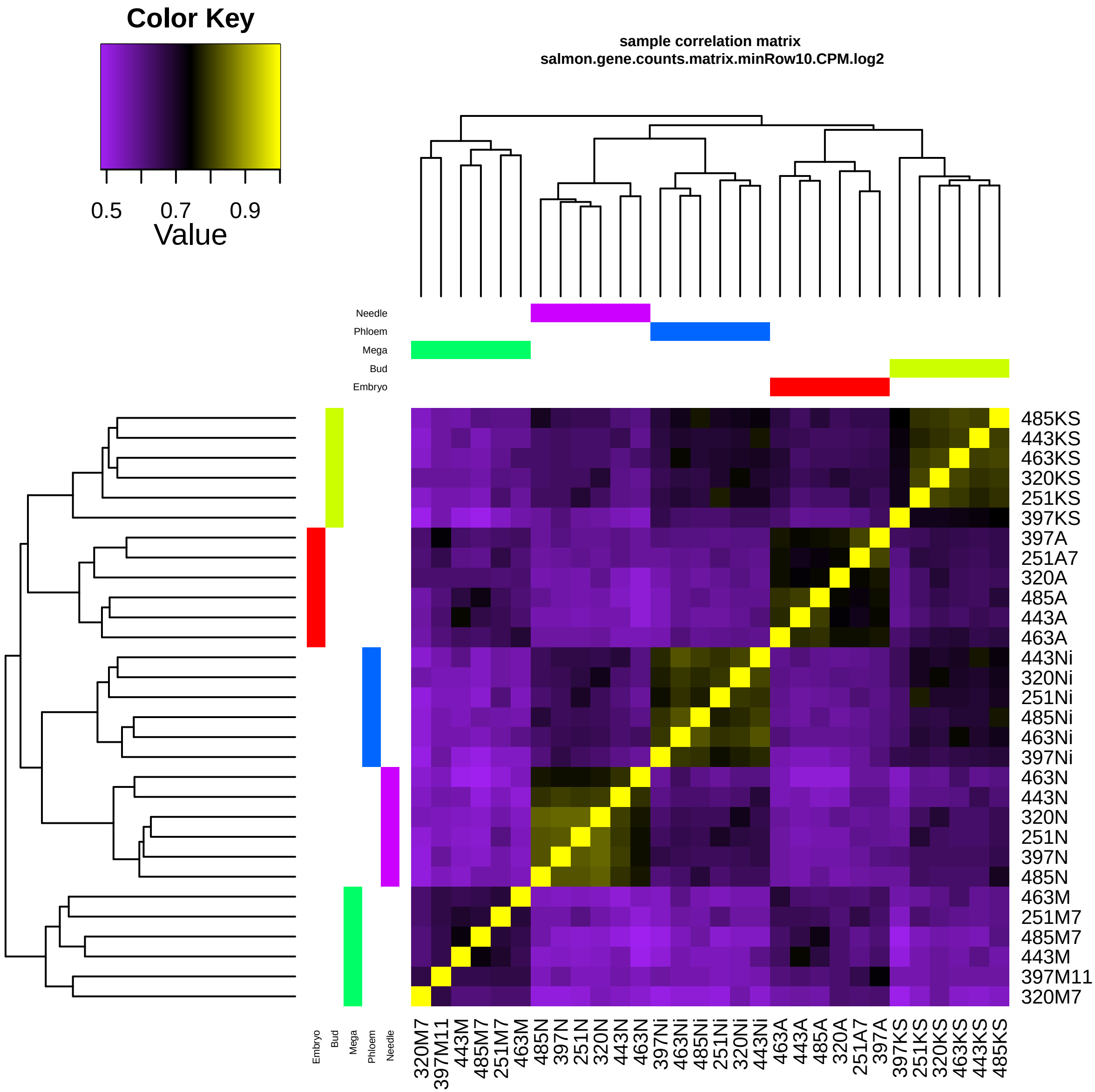
