## Supplementary material for "Atlas of tissue-specific and tissue-preferential gene expression in ecologically and economically significant conifer *Pinus sylvestris*": Tables supplementary files: Table S1. Tree number and coordinates.docx

| **Tree identifier** | **Tree location** | **Latitude** | **Longitude** |
| --- | --- | --- | --- |
| 215 | Ranta-Halola | 61.655 | 29.280 |
| 320 | Ranta-Halola | 61.657 | 29.292 |
| 397 | Mäkrä | 61.838 | 29.396 |
| 443 | Mäkrä | 61.838 | 29.393 |
| 463 | Mäkrä | 61.837 | 29.392 |
| 485 | Mäkrä | 61.836 | 29.394 |

**Table S1.** Tree identification number and location coordinates for the trees sampled at Punkaharju ISS.
